# Age-related differences in GABA: Impact of analysis technique

**DOI:** 10.1101/2022.02.21.481330

**Authors:** M. Simmonite, S.J. Peltier, T. A. Polk

**Author notes:** **CORRESPONDING AUTHOR:** Molly Simmonite, University of Michigan, Dept of Psychology, B033 East Hall, Ann Arbor, MI 48109.

## Abstract

Previous research using magnetic resonance spectroscopy (MRS) has indicated that GABA levels decline in multiple brain regions over the course of healthy aging. However, brain atrophy also occurs during healthy aging, and as a result the tissue composition of MRS voxels (i.e., the percentage of grey matter, white matter, and cerebrospinal fluid in the voxel) may also differ between age groups. Many authors therefore argue for applying a correction to GABA estimates in order to control for differences in tissue composition. Here, we use data from a large healthy aging study to investigate the influence of three tissue correction strategies on age-group differences in GABA. We also evaluate the use of different analysis packages and reference metabolites on group differences. A 3T MEGA-PRESS sequence was used to obtain spectra from seven voxels placed in the visual, auditory, and sensorimotor cortex of 58 young adults (aged 18-29 years) and 85 older adults. We obtained several different estimates of GABA concentrations from the spectra using two analysis software packages (Gannet 3.1 and LCModel), three reference metabolites (water, creatine and N-acetylaspartate) and four tissue correction strategies. Young adults consistently demonstrated significantly higher GABA concentrations in the visual, auditory, and sensorimotor cortex when we used an uncorrected GABA estimate referenced either to water or creatine. When uncorrected GABA was referenced to N-acetylaspartate, age-related differences were observed only in the right ventral visual cortex. Similarly, when any of the four tissue corrections were applied to the data, only age-related differences in the left and right ventral visual cortex voxels remained. Correlations between GABA concentration estimates obtained from different software packages were moderate, as were correlations between uncorrected GABA estimates when different baseline metabolites were used. Correlations between all tissue corrections we explored were extremely high. These results confirm that reports of age-related differences in GABA concentrations are driven, at least in part, by changes in tissue composition.

## 1. INTRODUCTION

Gamma-aminobutyric acid (GABA) is the primary inhibitory neurotransmitter in the human brain (McCormick, 1989), and plays a vital role in the function of the central nervous system. Previous research suggests that GABA levels decrease with age in a number of brain areas including frontal (Marenco et al., 2018; Porges, Woods, Edden, et al., 2017; Rowland et al., 2016), parietal (Gao et al., 2018; Porges, Woods, Edden, et al., 2017), temporal (Lalwani et al., 2019) and occipital (Chamberlain et al., 2021; Hermans et al., 2018; Simmonite et al., 2018) cortex. To date, no longitudinal investigations of GABA have been performed, however a recent meta-analysis of cross-sectional studies indicates a non-linear age-related trajectory—characterized by an increase in GABA in early life, stability during early adulthood and a gradual decrease in later life (Porges, Jensen, Foster, Edden, & Puts, 2021). Additionally, GABA levels are associated with clinical outcomes (Wenneberg et al., 2020) and behavioral performance in several domains, including visual function (Simmonite et al., 2018), motor control (Boy et al., 2010), tactile function (Puts, Edden, John Evans, McGlone, & McGonigle, 2011), sensory processing (Puts et al., 2017), and cognitive function (Porges, Woods, Edden, et al., 2017). It is important to note however, that other studies have found no significant age-related changes in GABA levels (Aufhaus et al., 2013; Mikkelsen et al., 2017) or even increased GABA levels in older adults (Pitchaimuthu et al., 2017).

The studies outlined above have utilized Magnetic Resonance Spectroscopy (MRS), a powerful technique which allows the direct detection of metabolites in the human brain non-invasively. While it can be challenging to isolate GABA signals since signals from other, more abundant metabolites can overlap and obscure GABA, recent advances in MRS techniques including the development of GABA-edited MEGA-PRESS (Mescher, Merkle, Kirsch, Garwood, & Gruetter, 1998) allow estimates of its concentration.

Brain atrophy is an important consideration for age-related neuroimaging research in general, and MR spectroscopy in particular. It is estimated that the adult brain shows a reduction of approximately 5% of brain weight each decade after the age of 40 years (Svennerholm, Boström, & Jungbjer, 1997). However, these changes are not uniform across the brain, but appear to occur on an anterior-posterior gradient with frontal and parietal regions demonstrating larger rates of decline when compared with temporal and occipital regions (Thambisetty et al., 2010). Furthermore, grey matter (GM), white matter (WM) and cerebrospinal fluid (CSF) exhibit different change patterns (Ge et al., 2002; Raz, 2004).

In light of age-related brain atrophy, it is important to consider the composition of MRS voxels when comparing metabolite concentrations between young and older participants, since both metabolite and reference signals differ between GM, WM and CSF (Harris, Puts, & Edden, 2015). Concentrations of GABA in grey matter are approximately double that of white matter, and negligible in CSF (Harris et al., 2015; Jensen, de B. Frederick, & Renshaw, 2005), and correcting for age differences in tissue composition between age-groups can therefore have an impact on age differences in GABA concentrations. Previous work by both Porges et al., (2017) and Maes et al (2018) explored GABA levels in voxels across the brain, applying several different correction strategies for the tissue composition of the voxel. These studies revealed that some age-related differences in GABA concentrations were not present when using a correction that accounted for the different amounts of GABA in grey and white matter— indicating that the selection of tissue correction can significantly impact the interpretation of MRS results.

The current study expands upon that work and explores the impact of different voxel tissue correction strategies in several cortical areas, in a sample comprised of healthy young (18 - 29) and older (65+) adults. We include several voxel locations in this manuscript—3 visual cortex voxels, 2 auditory voxels and 2 somatosensory voxels—and compare uncorrected GABA values alongside 4 tissue correction methods. We also investigate GABA concentrations obtained by different software packages, and referenced to different baseline metabolites, to more completely understand how analysis technique impacts age group differences. Additionally, we compare GABA levels across the brain to determine if GABA levels appear to be reduced uniformly across the brain during aging (i.e., do individuals with low visual GABA, also display low auditory and somatosensory GABA?).

The majority of MRS data from our sample has been previously reported (bilateral ventral visual voxels, Chamberlain et al. 2021); bilateral auditory cortex voxels, Lalwani et al. 2019); bilateral somatosensory voxels,(Cassady et al., 2019), and all of these studies found significant differences between the young and older groups. However, in each of these papers, values from bilateral voxels were combined to create a single GABA value per participant and only one or two different tissue corrections were considered. Furthermore, the relationship between GABA estimates in different brain areas was not explored.

## 2. METHODS AND MATERIALS

### 2.1. Participants

Participants were 58 young adults (mean age: 22.76 ± 2.86; 29 women and 29 men) and 85 older adults (mean age: 70.61 ± 5.10; 54 women and 31 men). All participants were healthy, right-handed, and were free of significant cognitive impairment (all participants had an NIH cognition score of > 85). Prior to taking part in the study, all participants were telephone screened to ensure that they were free from fMRI safety contraindications and psychotropic medications. Full inclusion/exclusion criteria can be found in Gagnon et al. (2019), which describes the protocol of the large, multi-modal project of which this dataset was part. All procedures were approved by the University of Michigan Institutional Review Board, and all participants provided written informed consent prior to taking part in the study.

### 2.2. MRS data acquisition

Scanning was performed on a GE Discovery MR750 3T scanner at the University of Michigan Functional Magnetic Resonance Imaging Laboratory. A high resolution T1-weighted spoiled 3D gradient-echo acquisition (SPGR) image was collected for MRS voxel placement and segmentation. GABA-edited MR spectra from seven voxels (left primary visual [LPV], bilateral ventral visual [LVV and RVV], bilateral auditory cortex [LAUD and RAUD] and bilateral sensorimotor [LSM and RSM]) were acquired using a MEGA-PRESS sequence with the following parameters: TE = 68 ms (TE1 = 15 ms, TE2 = 53 ms); TR =1800 ms; 256 transients (128 ON interleaved with 128 OFF) of 4096 data points; spectral width = 5 kHz; frequency selective editing pulses (14 ms) applied at 1.9 ppm (ON) and 7.46 ppm (OFF). Total scan time was approximately 8.5 min per voxel.

Voxel locations are shown in Figure 1. In each participant, the LPV voxel was placed according to a template image. The remaining six voxels were placed according to the individual’s region of maximal activity within an anatomical mask during a task-based fMRI session (data which are not presented in this manuscript). Full details are provided in Gagnon et al., (2019). For the ventral visual voxels, data from a faces/houses task was used during which participants were presented with blocks of face images and blocks of house images, as well as fixation blocks in which a fixation cross was presented. Contrast maps were computed for face blocks vs. fixation blocks and for house blocks vs. fixation blocks. Using these contrast maps, MRS voxels were placed to overlap with the areas of highest activation in ventral visual cortex for each hemisphere. For the auditory cortex voxels, data from an auditory task in which participants heard clips of instrumental music and foreign speech was used to place the MRS voxels, with music vs. no sound and foreign speech vs. no sound contrasts computed. The sensorimotor voxels were placed to maximize overlap with data from both a motor task in which participants were asked to perform button presses using their right and left thumbs, and a somatosensory task in which they received vibrotactile stimulation to fingers on their right and left hands. The LSM voxel was placed according to the right-hand activations (i.e., the right-hand vs no movement contrast map from the motor task and the right-hand vs no stimulation contrast map from the somatosensory task) and the RSM voxel was placed according to the left-hand activations (i.e., the left-hand vs no movement contrast map from the motor task and the left-hand vs no stimulation contrast map from the somatosensory task).

**Figure 1:**
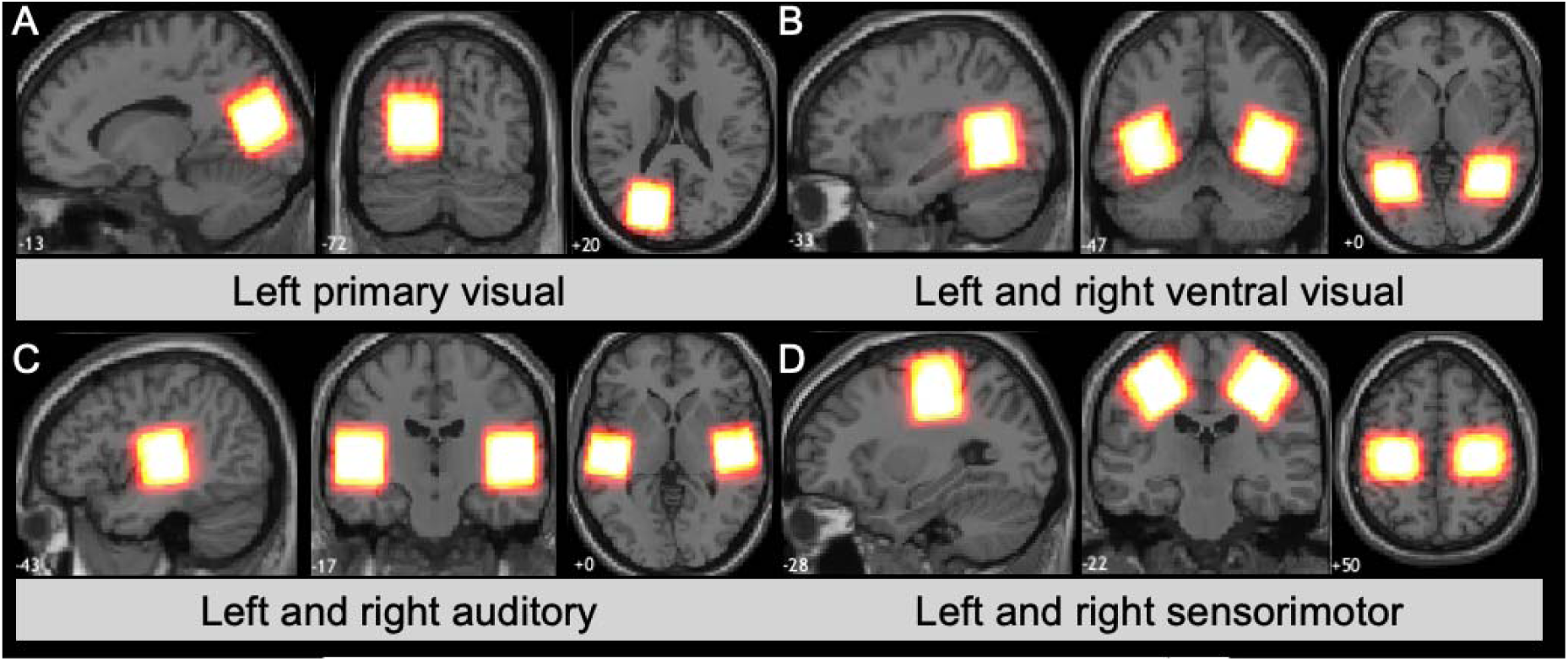
Voxel placement overlap for all participants, with brighter yellow color indicating greater overlap and red indicating less overlap. For bilateral voxel pairs, sagittal view shows the left voxel only. A. Left primary visual voxel (LPV) B. Left and right ventral visual voxel (LVV and RVV) C. Left and right auditory voxels (LAUD and RAUD) and D. Left and right sensorimotor voxels (LM and RSM)

### 2.3. MRS analysis

The Gannet 3.1 toolbox (Edden, Puts, Harris, Barker, & Evans, 2014) was used to preprocess and quantify the GABA signal from the seven voxels. Default parameters were used, including frequency and phase correction of the time-resolved data using spectral registration (Near et al., 2015). Since the resonance of macromolecules is also 3.0 ppm, co-edited MM signals overlap with GABA and make a significant (up to 50%) contribution to the GABA signal (Aufhaus et al., 2013). Therefore, all GABA values reported are GABA and macromolecules, and will be referred to as GABA+ in line with conventions in the literature. Since GABA+ concentrations in the ^1^H-MRS literature often differ with regards to the internal reference signal used, we include estimates of GABA+ referenced to the unsuppressed water signal (GABA+/H_2_O, as well as referenced to creatine (GABA+/Cr) and N-acetylaspartate (GABA+/NAA). LCModel (Provencher, 1993, 2001) was also used to quantify GABA+ concentrations from each of the seven voxels, to allow comparison between the two pieces of software.

Segmentation of the T1-weighted anatomical image was performed using the SPM12 segmentation function which is integrated into Gannet 3.1. This function allows the CSF, WM, and GM fractions of each voxel to be estimated for each participant. Four different tissue correction strategies are performed in Gannet, details of which are provided in Table 1.

**Table 1:** GABA+ estimation details

| Table 1: GABA+ estimation details |  |  |  |  |
| --- | --- | --- | --- | --- |
| Measure | Reference metabolite | Correction technique, | Software | Details |
| GABA+/H<sub>2</sub>O | Water | Uncorrected | Gannet 3.1 | Raw GABA+ concentrations with no correction applied, referenced to water |
| GABA+/Cr | Cr | Uncorrected | Gannet 3.1 | Raw GABA+ concentrations with no correction applied, creatine used as a reference metabolite. |
| GABA+/NAA | NAA | Uncorrected | Gannet 3.1 | Raw GABA+ concentrations with no correction applied, NAA used as a reference metabolite. |
| GABA+/H<sub>2</sub>O CSF | Water | CSF corrected | Gannet 3.1 | Raw GABA+ concentration divided by sum of voxel GM% and WM% |
| GABA+/H<sub>2</sub>O TC | Water | Tissue corrected | Gannet 3.1 | Like CSF correction, but also uses CSF%, WM% and GM% and controls for the differential relaxation constants and water visibility of each tissue type. |
| GABA+/H<sub>2</sub>O ATC | Water | α-tissue corrected | Gannet 3.1 | Like TC correction, but applies a weighting of .5 to the WM%, to account for the fact that GABA+ in GM is twice that in WM (this is analogous to correcting GABA to a voxel of 100% grey matter) |
| GABA+/H<sub>2</sub>O GATC | Water | Group α-tissue corrected | Gannet 3.1 | Like ATC correction but normalized to the average voxel tissue composition of the sample. In our case, we normalized to the average composition of the young participants. |
| GABA+/H<sub>2</sub>O LCM | Water | uncorrected | LCModel | As GABA+/H<sub>2</sub>O but estimated using LCModel software. |

Gannet 3.1 calculates a measure of fit error (the ratio of the standard deviation of the fit residual to the amplitude of the fitted peak). Gannet 3.1 derived GABA+ estimates were only included in analyses when the GABA+ fit error was below 15%. Similarly, LCModel estimates of GABA+ were only used in statistical analyses if the Cramer-Rao bounds were less than 20%. Details of these metrics are included in Supplementary Table 1.

### 2.4. Statistical analysis

Statistical analyses and visualizations were performed in R. To understand differences in voxel composition between the two age groups, two sample t-tests were performed for each of the tissue fractions (GM, WM and CSF), for each of the voxels (LPV, LVV, RVV, LAUD, RAUD, LSM and RSM). To investigate age-related differences in GABA+ concentrations, and how the choice of analysis software, reference metabolite, or correction strategy affect these differences, we performed two-sample t-tests comparing the GABA+ concentrations of young and older adults for each of the GABA+ estimates detailed in Table 1. Since we normalized all participants to the average young adult voxel, we did not report statistics on this measure as they are identical to those for GABA+/H_2_O ATC.

To determine whether GABA+ concentrations are uniform across the brain and if GABA+ concentrations decline uniformly across the brain, we calculated correlations between all possible pairs of the seven MRS voxels, first as partial correlations controlling for age in the whole sample and then in the two age groups separately. This was repeated for each of the GABA+ concentration estimates.

Finally, to investigate the impact of choice of software, reference metabolite or voxel tissue composition correction strategy, we calculated correlation coefficients between pairs of GABA+ estimates in the entire sample and in each age group separately. Here, we examined the impact of software by calculating the relationship between the GABA+/H_2_O estimate obtained in Gannet with the GABA+/H_2_O estimate obtained in LCModel; we investigated the impact of reference metabolite by calculating relationships between all possible pairs of the Gannet estimates of GABA+/H_2_O GABA+/Cr and GABA+/NAA; and we investigated the impact of voxel tissue composition correction strategy by calculating relationships between all possible pairs of the available Gannet corrections, GABA+/H_2_O (i.e., without a correction), GABA+/H_2_O CSF, GABA+/H_2_O TC, and GABA+/H_2_O ATC. Correlations were calculated for each of the seven voxels separately, and then averaged across the voxels. For the full participant sample, we calculated both bivariate correlations between the two variables of interest, as well as partial correlations which controlled for age.

## 3. RESULTS

### 3.1. Tissue Composition

Average voxel tissue compositions for each of the seven voxels are presented in Table 2. Across all seven voxels, older participants had greater CSF percentages (all *p*’s <.001) and younger participants had greater grey matter percentages (all *p*’s <.001). Age effects on white matter fraction were much smaller, but younger participants had greater white matter fractions in the LVV voxel and lower white matter fractions in LAUD, RAUD and LSM. There were no significant group differences between white matter percentages in LPV, RVV and LSM.

**Table 2:**
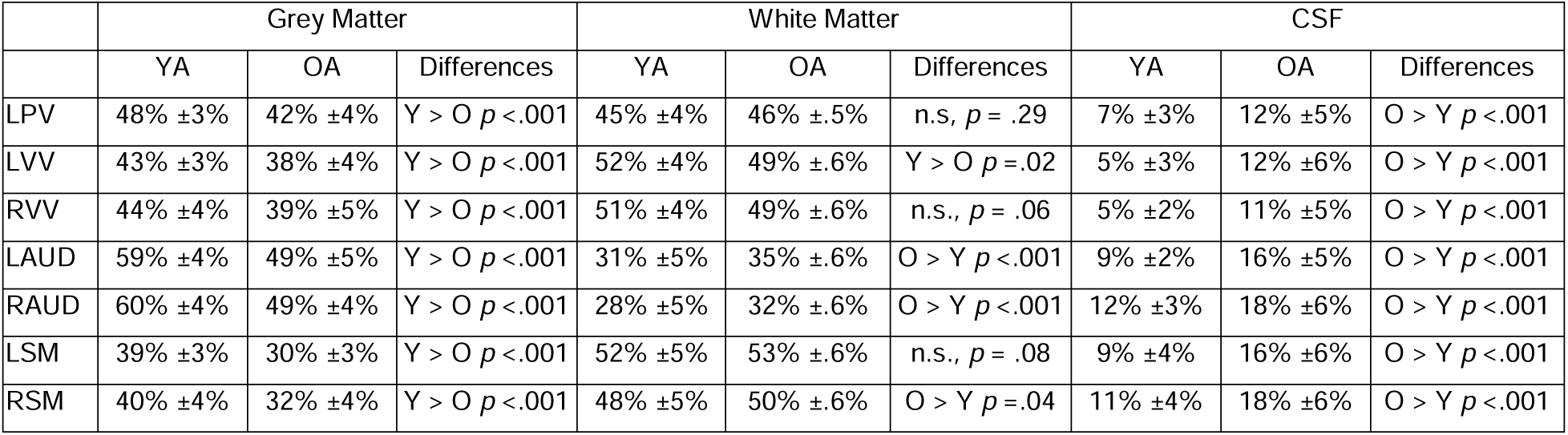
MRS voxel tissue compositions

### 3.2. Age differences in GABA+ concentrations

Age-related GABA+ differences are presented in Figure 2, and in Supplementary Tables 2,3 and 4. Young adults had significantly higher GABA+/H_2_O and GABA+/Cr than older adults across all seven voxels (all *p*s <.05). Young adults had higher GABA+/NAA in the RVV voxel (t(130)=2.53, p = .01), but there were no significant group differences in GABA+/NAA in the other voxels. Comparison of the corrected measures indicated left and right ventral visual GABA+/H_2_O CSF, GABA+/H_2_O TC and GABA+/H_2_O ATC concentrations were higher in the young adults compared with the older adults (all *p*’s < .01) but there were no differences in the remaining voxels. Analysis of LCModel estimates of GABA+/H_2_O, revealed significantly higher concentrations in the left and right ventral visual, auditory, and sensorimotor cortex voxels (all *p*’s <.005) but differences did not reach significance in the LPV voxel (t(131) = 1.90, p = .06).

**Figure 2:**
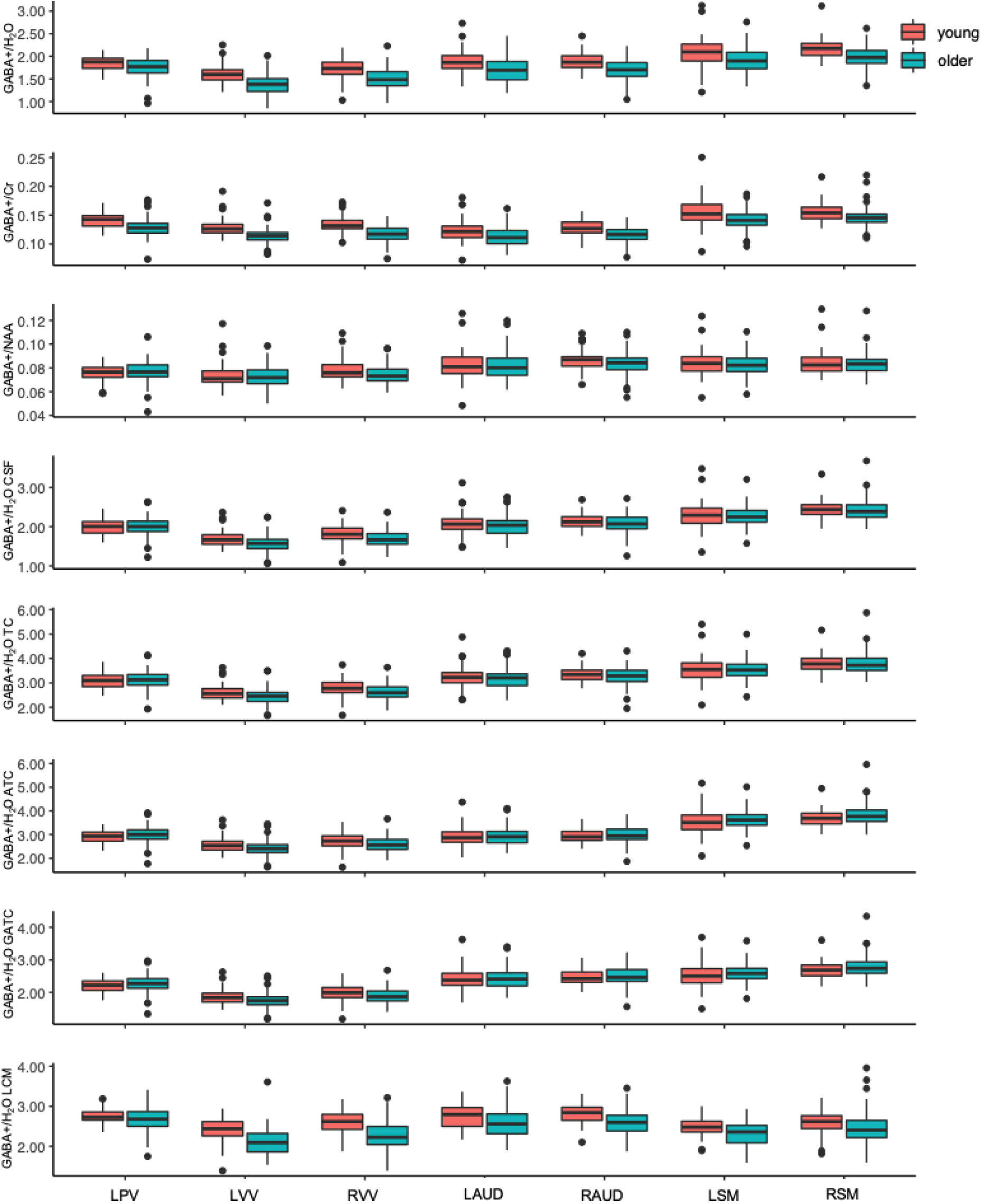
Group differences in GABA+ concentration estimates for left primary visual cortex (LPV), left and right ventral visual cortex (LVV and RVV), left and right auditory (LAUD and RAUD), and left and right sensorimotor cortex (LSM and RSM) voxels. Older adults are presented in blue and young adults are presented in red. Top row presents uncorrected GABA+/H_2_O concentrations estimated in Gannet, second row presents GABA+/Cr concentrations, third row presents GABA+/NAA concentrations, fourth row presents GABA+/H_2_O concentrations after CSF correction in Gannet 3.1, fifth row presents GABA+/H_2_O concentrations following tissue correction (TC) in Gannet 3.1, sixth row presents GABA+/H_2_O concentrations following alpha-tissue correction (ATC) in Gannet 3.1, seventh row presents GABA+/H_2_O concentrations after group alpha-tissue correction (GATC) in Gannet 3.1 and bottom row presents GABA+/H_2_O estimated in LCModel.

### 3.3. Correlations between GABA+ estimates in different brain regions

Correlations between GABA+ concentrations are presented in Figure 3, with correlations which met a significance threshold of p <.05 (uncorrected for multiple comparisons) highlighted with a black border. Those correlations that remained significant following correction for multiple comparisons are highlighted with a yellow border.

**Figure 3:**
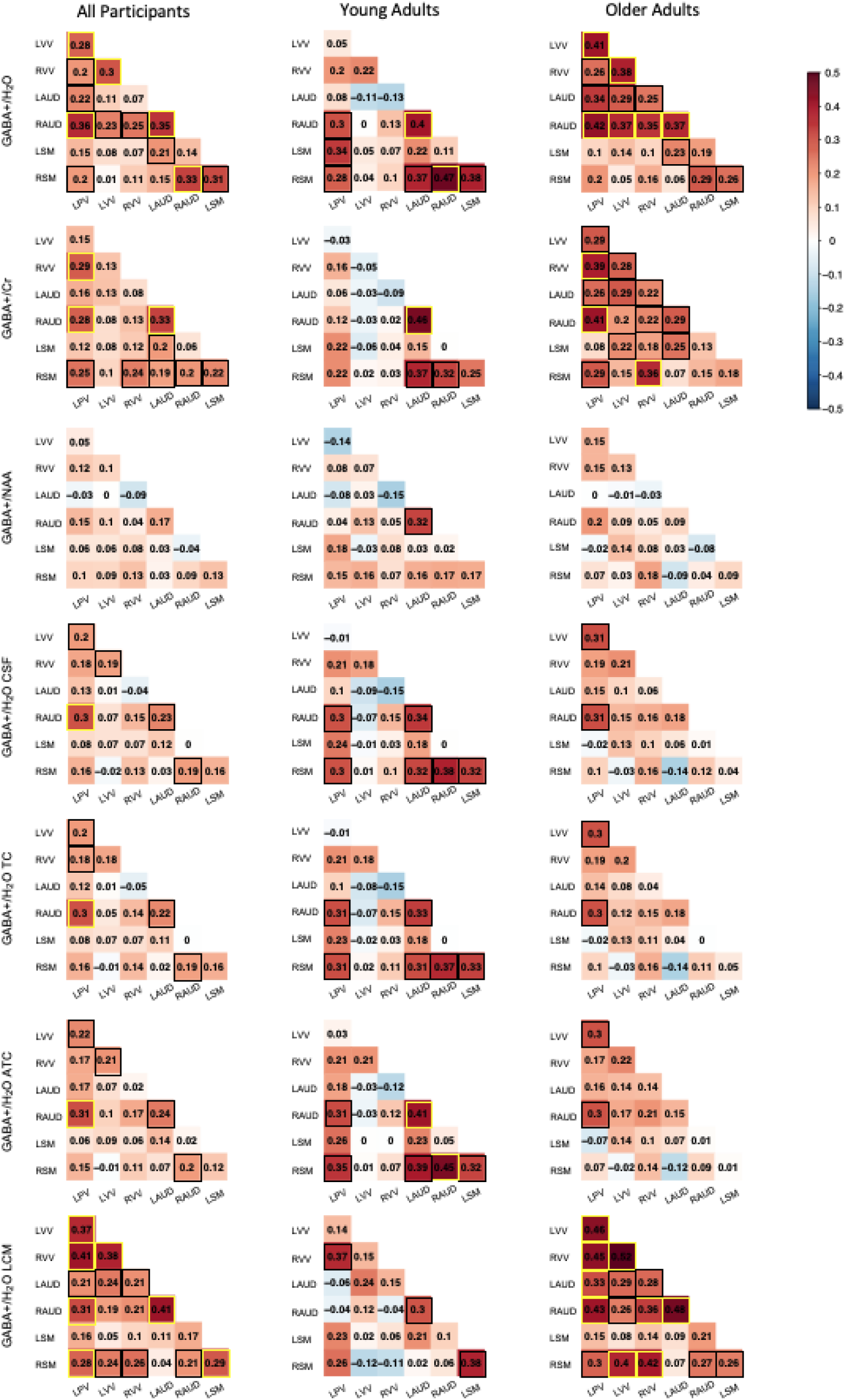
Matrices showing correlations between GABA+ concentration estimates in each of the seven voxels. First column shows partial correlations in the whole sample, while controlling for age, second column shows correlations in the young sample only and third column shows correlations in the older sample. Box color indicates the strength of the correlation, which is also presented in black text. Boxes with a black background represent correlations which are significant at p<.05 uncorrected, and boxes with a yellow background denote correlations that survive correction for multiple comparisons.

When we looked at relationships between GABA+/H_2_O across the different voxels, we found broadly similar patterns of relationships between voxels across all participants and in the older group. There were significant correlations between LVV and RVV, LPV and LVV, LAUD and RAUD, and LPV and RAUD, which survived correction for multiple comparisons. There were fewer significant relationships between GABA+/H_2_O concentrations in different voxels in the young adults, but the correlation between LAUD and RAUD, and the correlation between RSM and RAUD survived correction for multiple comparisons.

Patterns of relationships across the different tissue corrections appeared broadly similar to one another. Across all participants and in the older adults, relationships between LPV and LVV, and LPV and RAUD were the strongest. The relationship between LPV and RAUD was also present in the young participants, who also demonstrated significant relationships between LAUD and RAUD, and the RSM voxel and the LAUD, RAUD and LSM voxels.

For GABA+/Cr, we saw correlations between LAUD and RAUD surviving corrections for multiple comparisons across all participants, as well as in the young participants. Correlations between LPV and both RVV and RAUD survived correction across all participants and in the older participant group. For GABA+/NAA, there were few significant correlations between voxels, and none that remained significant following correction for multiple comparisons, whether we examined the entire sample or just the young or older participants.

Analysis of LCModel estimates of GABA+/H_2_O revealed that across all participants and in the older group, GABA+/H_2_O concentrations correlated significantly across most voxel pairs, with notable exception of pairs which included the LSM voxel. Additionally, correlations between GABA+/H_2_O concentrations in the LPV voxel and the ventral visual, auditory and right sensorimotor voxels appeared particularly robust, as did correlations between left and right ventral visual voxels, between left and right auditory voxels and between left and right sensorimotor voxels

### 3.4. Correlations between different GABA+ estimates

Figure 4 presents correlations between the different GABA+ estimates, for the whole sample (both with and without controlling for age), for young participants and for older participants. Across the seven voxels, average correlations between the estimates of GABA+/H_2_O across the whole sample were r = .52 ±.17 when bivariate correlations were calculated, and r = .52 ±.08 when we controlled for age. These correlations were r = .36 ±.15 in the young group, and r = .53±.19 in the older group. When we compared the use of three different reference metabolites (H_2_O, Cr and NAA), we found bivariate correlations across the whole sample ranging from r = .58 ±.21 to r = .69 ±.25, and correlations ranging from .70 ±.09 to .81 ±.05 when we controlled for age. In the young group, these correlations ranged from .60 ±.24 to r = .73 ±.27, and in the older group they ranged from r = .59 ±.23 to .72 ±.26. Across all participants, correlations between GABA+/H_2_O. (i.e. the uncorrected GABA+ measure obtained from Gannet) and the three Gannet corrections (CSF, TC and ATC) ranged from r = .83 ±.09 to r = .89 ±.06 when bivariate correlations were calculated and r = .89 ±.06 to r = .92 ±.05 when we controlled for age. These correlations ranged from r = .94 ±.05 to r = .95 ±.04 in the young adults and r = .86 ±.06 to r = .89 ±.05 in the older adults. Comparison of the three corrections revealed extremely strong relationships, ranging from r = .97 ±.01 to 1.00 ±.00 across all participants, and in both the young and older adults.

**Figure 4:**
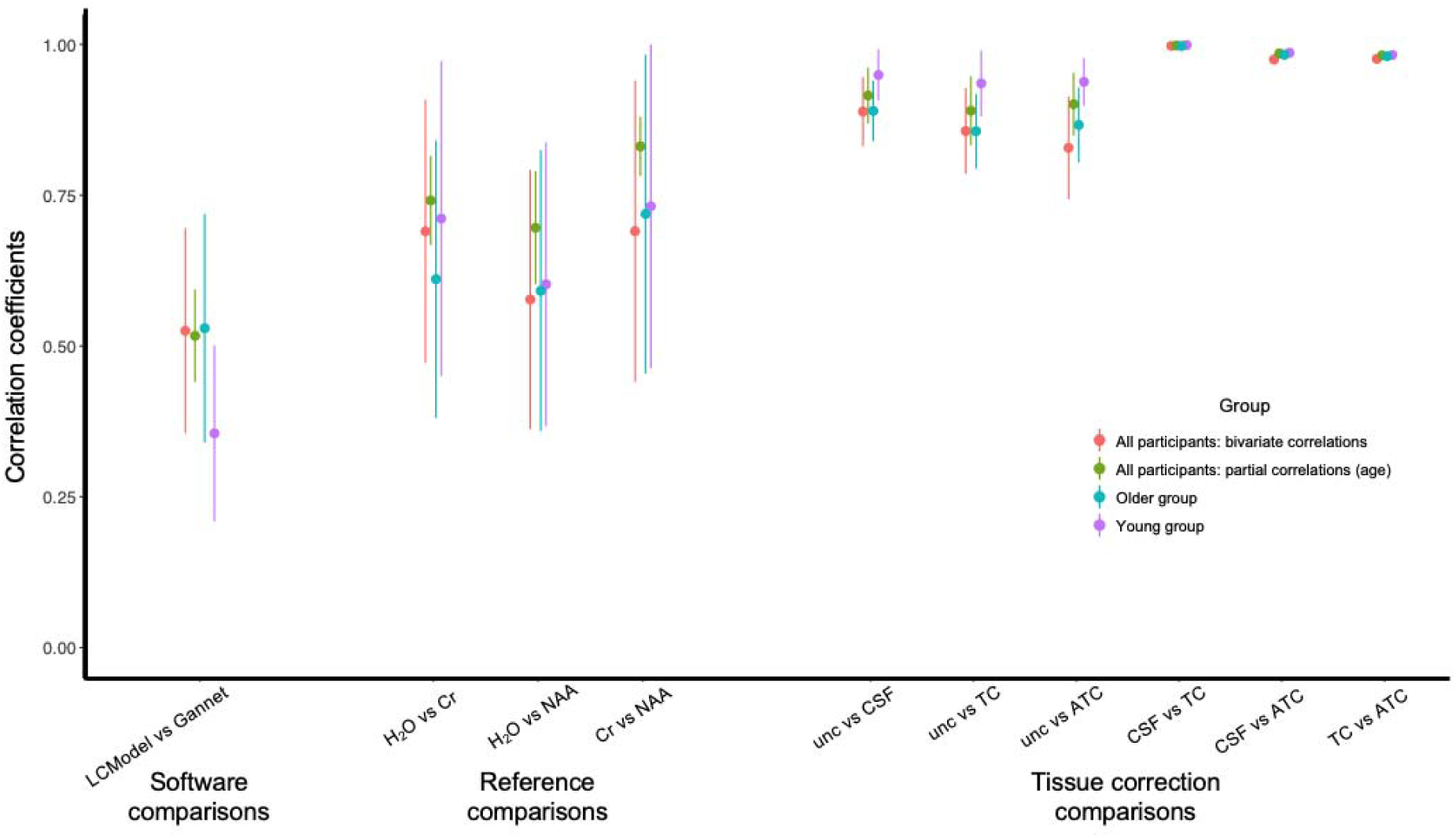
Average correlations between different GABA+ estimates. Bivariate correlations calculated for the whole sample are presented in red, partial correlations across the whole sample controlling for age are presented in green, older adults are presented in blue and young adults are presented in purple. Filled circles denote average correlations across the 7 MRS voxels, with lines extending out to show standard deviations. ATC, alpha-tissue correction; Cr, creatine; CSF, cerebrospinal fluid correction; H_2_O, water; NAA, n-Acetyl aspartate; TC, tissue correction; unc, uncorrected.

## 4. Discussion

The present study examined the impact of tissue correction, reference metabolite and software choice on age-related group differences in GABA+ concentration. We observed changes in tissue composition of the brain over healthy aging, and that visual, auditory, and sensorimotor cortex, age-related GABA+ reductions were dependent upon which reference metabolite was used, and whether the GABA+ concentration estimate was corrected for the tissue composition of the voxel. These variables also influenced the relationships between GABA concentrations across the brain.

### 4.1. Tissue correction of water-referenced GABA

We applied four different tissue corrections to our dataset. The first, and most basic, was the CSF correction, in which it is assumed that the amount of GABA present in CSF is negligible, and thus we divide the uncorrected GABA measurement by the fraction of the voxel that is made up of GM or WM. More sophisticated is the TC, which similarly corrects for the fraction of CSF in the voxel, but also accounts for differences in the visibility and relaxation of the water signals in each of the tissue types. The alpha-tissue correction builds on TC and accounts for the different amounts of GABA in grey and white matter, with the applied alpha of 0.5 being the assumed ratio of GABA+ in WM to GM. Our final correction, GATC then normalises these voxels to a group-average tissue composition, allowing the comparison between individuals. Since we demonstrated age-group differences in the tissue compassion of all seven voxels, normalising each age group to its group average tissue composition would introduce a bias, and so we chose to normalise both groups to the average young adult voxels.

Consistent with previous reports (Maes et al., 2018; Porges, Woods, Lamb, et al., 2017), we found that the type of tissue correction used has an impact on age-related findings. We found group differences in the visual, auditory, and sensorimotor cortices for uncorrected GABA+ estimates, but only ventral visual group differences remained after any type of correction. We also found extremely high correlations between our corrected GABA+ concentration estimates. Taken together, these results suggest that what makes the greatest difference is the application (or not) of tissue correction, rather than the precise type of correction applied.

However, it should be noted that when we used a correction for multiple comparisons across the voxels, this slightly changed the pattern of the results. Here, only age-group differences for the right ventral visual voxel remain significant when CSF correction or TC are used, and no group differences are significant when an ATC correction was used. This demonstrates that there are subtle differences between the types of tissue correction. Further, when we look at the average GABA+/H_2_O ATC concentrations, we see that older adults have higher values than young adults in the LPV, LSM and RSM voxels, although these differences are not significant (p’s = .13 to .17). This observation is consistent with findings by Maes et al., (2018) who found similar trends in the occipital cortex.

### 4.2. Use of Creatine or NAA as a reference metabolite

As well as assessing age-related differences in GABA+ referenced to water, we also examined GABA+ concentrations when referenced to Cr and NAA. We found that Cr-referenced GABA+ was lower in older adults across all seven voxels, whereas when NAA was used as a reference metabolite, there were only age-related differences observed in the RVV, and these differences did not survive correction for multiple comparisons. While we found largely comparable GABA+/NAA concentrations between the age groups in the remaining six voxels, Maes et al., (2018) compared these reference compounds and found increased GABA+/NAA in older adults. Our results are more consistent with Gao et al., (2018) who saw age-related declines in both GABA+/Cr and GABA+/NAA in frontal and parietal voxels.

Use of Cr or NAA as reference metabolites are impacted by how these metabolites themselves decline over age. A systematic review, performed by Cleeland and colleagues (2019) suggests that NAA declines with age, but presents more mixed findings for Cr. While no reference signal is perfect, with each having their own merits (Mullins et al., 2014), caution should be used when evaluating age-related GABA+ differences when Cr or NAA metabolites are used as references.

### 4.3. Comparison of uncorrected GABA+/H_2_O estimates obtained by Gannet 3.1 and LCModel

We analysed the MRS data we obtained using two highly popular pieces of software, Gannet 3.1 and LCModel. These packages take different approaches to modelling spectra. LCModel estimates GABA voxels using a linear combination modelling approach, using a basis set of metabolites. Gannet, on the other hand, was specifically developed for the analysis of GABA-edited MRS data, and models GABA by fitting a single Gaussian peak.

Although we found a similar pattern of age-related group differences for GABA+/H_2_O (significant differences across the ventral visual, auditory, and sensorimotor voxels which survive correction for multiple comparisons), we saw only moderate correlations between the estimates for both young and older adults. Additionally, the absolute values of GABA+/H_2_O estimates were quite different, ranging from 1.60 - 2.17 in young adults and 1.38 - 1.98 in older adults when estimated using Gannet 3.1, and 2.41 - 2.81 in young adults and 2.11 to 2.67 in older adults when estimated using LCModel. These findings are in line with those of Craven et al., (2021) who performed a large-scale evaluation of anterior cingulate cortex MRS data obtained from 222 healthy adults aged 18 - 35 across 20 research sites, which was analysed using seven different algorithms. This study also found moderate correlation (.4) between estimates obtained using Gannet and LCModel, and GABA values obtained from LCModel appeared higher than those obtained from Gannet.

### 4.4. Relationships between GABA+ concentration estimates in different brain regions

Not only did the choice of reference metabolite, analysis software and voxel tissue correction strategy impact age-related GABA+ differences, but it also influenced relationships between different voxels. Prior evidence has indicated that, while there are significant correlations between GABA+ concentrations in bilateral homologous brain regions (Puts et al., 2018), GABA+ concentrations do not significantly correlate between brain regions (Boy et al., 2010; Grachev & Apkarian, 2000; Grachev, Swarnkar, Szeverenyi, Ramachandran, & Vania Apkarian, 2001; Greenhouse, Noah, Maddock, & Ivry, 2016) although inter-regional correlations do increase in middle age (Grachev et al., 2001).

Across all participants, we saw correlations between GABA+ concentrations in left and right visual, auditory, and somatosensory voxels, although correlations between left and right somatosensory GABA+ were only significant in young adults when the estimate was corrected for the tissue composition of the voxel. In older adults, uncorrected GABA+/H_2_O concentrations were significantly correlated in different brain regions, however none of these relationships survived correction for multiple comparison in measures of GABA+ which were corrected for tissue composition, suggesting that, rather than being the result of an age-related global alteration in GABA+ concentrations, these relationships were instead driven by tissue changes.

### 4.5. Limitations

The current study included a cross-sectional sample of young and older adults, which means that we cannot exclude the possibility that any group differences observed were the result of cohort effects—differences between the two groups (e.g., in education, nutrition, or life experiences) that covary with, but are unrelated to, age. The current study also only included young and older participants, with no middle-aged group, which meant we were unable to assess GABA+ concentrations and tissue changes across the life span. Finally, while MRS is an extremely useful way of measuring GABA+ in vivo, it only estimates GABA concentration, not inhibitory activity, and it does not differentiate between intra- and extra-cellular GABA. It is therefore difficult to draw inferences about the neural mechanisms underlying the observed age differences.

### 4.6. Conclusions

The results of our study indicate that healthy aging is accompanied by a decline in GABA+ concentrations, and that these declines are influenced by the changes in tissue composition that occur over the aging process. While we saw reduced uncorrected GABA+ concentrations in the visual, auditory, and sensory cortices, only right ventral visual differences remained after correction for voxel tissue composition, suggesting that there are other factors involved. Since we found only moderate correlations between GABA+ estimates using different software and reference metabolites, despite being computed from the same data, care should be taken when comparing studies.

## References

Aufhaus, E., Weber-Fahr, W., Sack, M., Tunc-Skarka, N., Oberthuer, G., Hoerst, M., … Ende, G. (2013). Absence of changes in GABA concentrations with age and gender in the human anterior cingulate cortex: A MEGA-PRESS study with symmetric editing pulse frequencies for macromolecule suppression. Magnetic Resonance in Medicine, 69(2), 317–320. https://doi.org/10.1002/mrm.24257

Boy, F., Evans, C. J., Edden, R. A. E., Singh, K. D., Husain, M., & Sumner, P. (2010). Individual differences in subconscious motor control predicted by GABA concentration in SMA. Current Biology, 20(19), 1779–1785. https://doi.org/10.1016/j.cub.2010.09.003

Cassady, K., Gagnon, H., Lalwani, P., Simmonite, M., Foerster, B., Park, D. C., … Polk, T. A. (2019). Sensorimotor network segregation declines with age and is linked to GABA and to sensorimotor performance. NeuroImage, 186, 234–244. Retrieved from https://www.sciencedirect.com/science/article/pii/S1053811918320780

Chamberlain, J. D., Gagnon, H., Lalwani, P., Cassady, K., Simmonite, M., Seidler, R. D., … Polk, T. A. (2021). GABA levels in ventral visual cortex decline with age and are associated with neural distinctiveness. Neurobiology of Aging, 102, 170–177. https://doi.org/10.1016/j.neurobiolaging.2021.02.013

Cleeland, C., Pipingas, A., Scholey, A., & White, D. (2019, March 1). Neurochemical changes in the aging brain: A systematic review. Neuroscience and Biobehavioral Reviews. Pergamon. https://doi.org/10.1016/j.neubiorev.2019.01.003

Craven, A. R., Bhattacharyya, P. K., Clarke, W. T., Dydak, U., Edden, R. A. E., Ersland, L., … Oeltzschner, G. (2021). Comparison of seven modelling algorithms for GABA-edited 1H-MRS. BioRxiv. https://doi.org/10.1101/2021.11.15.468534

Edden, R. A. E., Puts, N. A. J., Harris, A. D., Barker, P. B., & Evans, C. J. (2014). Gannet: A batch-processing tool for the quantitative analysis of gamma-aminobutyric acid-edited MR spectroscopy spectra. Journal of Magnetic Resonance Imaging, 40(6), 1445–1452. https://doi.org/10.1002/jmri.24478

Gagnon, H., Simmonite, M., Cassady, K., Chamberlain, J., Freiburger, E., Lalwani, P., … Polk, T. A. (2019). Michigan Neural Distinctiveness (MiND) study protocol: investigating the scope, causes, and consequences of age-related neural dedifferentiation. BMC Neurology, 19(1), 61. https://doi.org/10.1186/s12883-019-1294-6

Gao, F., Yin, X., Edden, R. A., Evans, A. C., Xu, J., Cao, G., … Wang, G. (2018). Altered hippocampal GABA and glutamate levels and uncoupling from functional connectivity in multiple sclerosis. Hippocampus, 28(11), 813–823. https://doi.org/10.1002/hipo.23001

Ge, Y., Grossman, R. I., Babb, J. S., Rabin, M. L., Mannon, L. J., & Kolson, D. L. (2002). Age-related total gray matter and white matter changes in normal adult brain. Part I: Volumetric MR imaging analysis. American Journal of Neuroradiology, 23(8), 1327–1333. Retrieved from http://www.ajnr.org/content/23/8/1327.short

Grachev, I. D., & Apkarian, A. V. (2000). Chemical heterogeneity of the living human brain: A proton MR spectroscopy study on the effects of sex, age, and brain region. NeuroImage, 11(5 I), 554–563. https://doi.org/10.1006/nimg.2000.0557

Grachev, I. D., Swarnkar, A., Szeverenyi, N. M., Ramachandran, T. S., & Vania Apkarian, A. (2001). Aging alters the multichemical networking profile of the human brain: An in vivo 1H-MRS study of young versus middle-aged subjects. Journal of Neurochemistry, 77(1), 292–303. https://doi.org/10.1046/j.1471-4159.2001.00238.x

Greenhouse, I., Noah, S., Maddock, R. J., & Ivry, R. B. (2016). Individual differences in GABA content are reliable but are not uniform across the human cortex. NeuroImage, 139, 1–7. https://doi.org/10.1016/j.neuroimage.2016.06.007

Harris, A. D., Puts, N. A. J., & Edden, R. A. E. (2015). Tissue correction for GABA-edited MRS: Considerations of voxel composition, tissue segmentation, and tissue relaxations. Journal of Magnetic Resonance Imaging, 42(5), 1431–1440. https://doi.org/10.1002/jmri.24903

Hermans, L., Leunissen, I., Pauwels, L., Cuypers, K., Peeters, R., Puts, N. A. J., … Swinnen, S. P. (2018). Brain GABA levels are associated with inhibitory control deficits in older adults. Journal of Neuroscience, 38(36), 7844–7851. https://doi.org/10.1523/JNEUROSCI.0760-18.2018

Jensen, J. E., de B. Frederick, B., & Renshaw, P. F. (2005). Grey and white matter GABA level differences in the human brain using two-dimensional, J-resolved spectroscopic imaging. NMR in Biomedicine, 18(8), 570–576. https://doi.org/10.1002/nbm.994

Lalwani, P., Gagnon, H., Cassady, K., Simmonite, M., Peltier, S., Seidler, R. D., … Polk, T. A. (2019). Neural distinctiveness declines with age in auditory cortex and is associated with auditory GABA levels. NeuroImage, 201. https://doi.org/10.1016/j.neuroimage.2019.116033

Maes, C., Hermans, L., Pauwels, L., Chalavi, S., Leunissen, I., Levin, O., … Swinnen, S. P. (2018). Age-related differences in GABA levels are driven by bulk tissue changes. Human Brain Mapping, 39(9), 3652–3662. https://doi.org/10.1002/hbm.24201

Marenco, S., Meyer, C., van der Veen, J. W., Zhang, Y., Kelly, R., Shen, J., … Berman, K. F. (2018). Role of gamma-amino-butyric acid in the dorsal anterior cingulate in age-associated changes in cognition. Neuropsychopharmacology, 43(11), 2285– 2291. https://doi.org/10.1038/s41386-018-0134-5

McCormick, D. A. (1989). GABA as an inhibitory neurotransmitter in human cerebral cortex. Journal of Neurophysiology, 62(5), 1018–1027. https://doi.org/10.1152/jn.1989.62.5.1018

Mescher, M., Merkle, H., Kirsch, J., Garwood, M., & Gruetter, R. (1998). Simultaneous in vivo spectral editing and water suppression. NMR in Biomedicine, 11(6), 266– 272. https://doi.org/10.1002/(SICI)1099-1492(199810)11:6<266::AID-NBM530>3.0.CO;2-J

Mikkelsen, M., Barker, P. B., Bhattacharyya, P. K., Brix, M. K., Buur, P. F., Cecil, K. M., … Edden, R. A. E. (2017). Big GABA: Edited MR spectroscopy at 24 research sites. NeuroImage, 159, 32–45. https://doi.org/10.1016/j.neuroimage.2017.07.021

Mullins, P. G., McGonigle, D. J., O’Gorman, R. L., Puts, N. A. J., Vidyasagar, R., Evans, C. J., … Wilson, M. (2014, February 1). Current practice in the use of MEGA-PRESS spectroscopy for the detection of GABA. NeuroImage. Academic Press. https://doi.org/10.1016/j.neuroimage.2012.12.004

Near, J., Edden, R., Evans, C. J., Paquin, R., Harris, A., & Jezzard, P. (2015). Frequency and phase drift correction of magnetic resonance spectroscopy data by spectral registration in the time domain. Magnetic Resonance in Medicine, 73(1), 44–50. https://doi.org/10.1002/mrm.25094

Pitchaimuthu, K., Wu, Q. Z., Carter, O., Nguyen, B. N., Ahn, S., Egan, G. F., & McKendrick, A. M. (2017). Occipital GABA levels in older adults and their relationship to visual perceptual suppression. Scientific Reports, 7(1). https://doi.org/10.1038/s41598-017-14577-5

Porges, E. C., Jensen, G., Foster, B., Edden, R. A. E., & Puts, N. A. J. (2021). The trajectory of cortical gaba across the lifespan, an individual participant data meta-analysis of edited mrs studies. ELife, 10. https://doi.org/10.7554/eLife.62575

Porges, E. C., Woods, A. J., Edden, R. A. E., Puts, N. A. J., Harris, A. D., Chen, H., … Cohen, R. A. (2017). Frontal Gamma-Aminobutyric Acid Concentrations Are Associated With Cognitive Performance in Older Adults. Biological Psychiatry: Cognitive Neuroscience and Neuroimaging, 2(1), 38–44. https://doi.org/10.1016/j.bpsc.2016.06.004

Porges, E. C., Woods, A. J., Lamb, D. G., Williamson, J. B., Cohen, R. A., Edden, R. A. E., & Harris, A. D. (2017). Impact of tissue correction strategy on GABA-edited MRS findings. NeuroImage, 162, 249–256. https://doi.org/10.1016/j.neuroimage.2017.08.073

Provencher, S. W. (1993). Estimation of metabolite concentrations from localized in vivo proton NMR spectra. Magnetic Resonance in Medicine, 30(6), 672–679. https://doi.org/10.1002/mrm.1910300604

Provencher, S. W. (2001). Automatic quantitation of localized in vivo 1H spectra with LCModel. NMR in Biomedicine, 14(4), 260–264. https://doi.org/10.1002/nbm.698

Puts, N. A. J., Edden, R. A. E., John Evans, C., McGlone, F., & McGonigle, D. J. (2011). Regionally specific human GABA concentration correlates with tactile discrimination thresholds. Journal of Neuroscience, 31(46), 16556–16560. https://doi.org/10.1523/JNEUROSCI.4489-11.2011

Puts, N. A. J., Heba, S., Harris, A. D., Evans, C. J., McGonigle, D. J., Tegenthoff, M., … Edden, R. A. E. (2018). GABA levels in left and right sensorimotor cortex correlate across individuals. Biomedicines, 6(3). https://doi.org/10.3390/biomedicines6030080

Puts, N. A. J., Wodka, E. L., Harris, A. D., Crocetti, D., Tommerdahl, M., Mostofsky, S. H., & Edden, R. A. E. (2017). Reduced GABA and altered somatosensory function in children with autism spectrum disorder. Autism Research, 10(4), 608–619. https://doi.org/10.1002/AUR.1691

Raz, N. (2004). The Aging Brain Observed in Vivo Differential Changes and Their Modifiers. In Cognitive Neuroscience of Aging: Linking Cognitive and Cerebral Aging (pp. 17–55). Retrieved from https://psycnet.apa.org/record/2004-21799-002

Rowland, L. M., Krause, B. W., Wijtenburg, S. A., McMahon, R. P., Chiappelli, J., Nugent, K. L., … Hong, L. E. (2016). Medial frontal GABA is lower in older schizophrenia: A MEGA-PRESS with macromolecule suppression study. Molecular Psychiatry, 21(2), 198–204. https://doi.org/10.1038/mp.2015.34

Simmonite, M., Carp, J., Foerster, B. R., Ossher, L., Petrou, M., Weissman, D. H., & Polk, T. A. (2018). Age-Related Declines in Occipital GABA are Associated with Reduced Fluid Processing Ability. Academic Radiology, 26(8), 1053–1061. https://doi.org/10.1016/j.acra.2018.07.024

Svennerholm, L., Boström, K., & Jungbjer, B. (1997). Changes in weight and compositions of major membrane components of human brain during the span of adult human life of Swedes. Acta Neuropathologica, 94(4), 345–352. https://doi.org/10.1007/s004010050717

Thambisetty, M., Wan, J., Carass, A., An, Y., Prince, J. L., & Resnick, S. M. (2010). Longitudinal changes in cortical thickness associated with normal aging. NeuroImage, 52(4), 1215–1223. https://doi.org/10.1016/j.neuroimage.2010.04.258

Wenneberg, C., Glenthøj, B. Y., Glenthøj, L. B., Fagerlund, B., Krakauer, K., Kristensen, T. D., … Nordentoft, M. (2020). Baseline measures of cerebral glutamate and GABA levels in individuals at ultrahigh risk for psychosis: Implications for clinical outcome after 12 months. European Psychiatry, 63(1). https://doi.org/10.1192/j.eurpsy.2020.77

