## Supplemental Tables for "Age-related differences in GABA: Impact of analysis technique"

| Supplementary Table 1: MRS quality metrics. | | | | | | | | |
| --- | --- | --- | --- | --- | --- | --- | --- | --- |
|  | Gannet 3.1 Fit Error (%) | | | | LCModel Cramér-Rao Lower Bounds (%SD) | | | |
|  | Mean ± SD | | t | *p* | Mean ± SD | | t | *p* |
|  | Young  adults | Older  adults |  |  | Young adults | Older  adults |  |  |
| LPV | 3.35 ±0.67 | 3.17 ±0.75 | 1.38 | .17 | 5.81 ±0.75 | 5.30 ±0.78 | 3.78 | .0002 |
| LVV | 4.32 ±1.52 | 4.25 ±1.25 | 0.29 | .77 | 6.11 ±0.76 | 5.90 ±0.74 | 1.62 | .11 |
| RVV | 4.14 ±1.51 | 3.58 ±0.86 | 2.81 | .006 | 5.76 ±0.79 | 5.77 ±0.73 | -0.01 | .99 |
| LAUD | 4.38 ±1.60 | 4.42 ±1.38 | -0.12 | .90 | 6.07 ±1.12 | 5.74 ±0.72 | 2.10 | .04 |
| RAUD | 3.64 ±1.05 | 3.93 ±1.19 | -1.47 | .15 | 5.77 ±0.73 | 5.75 ±0.66 | 0.16 | .88 |
| LSM | 4.78 ±1.86 | 4.36 ±1.30 | -1.58 | .12 | 6.37 ±0.84 | 6.42 ±1.11 | -.32 | .75 |
| RSM | 3.91 ±1.65 | 3.88 ±1.23 | 0.12 | .91 | 6.31 ±0.86 | 6.06 ±1.17 | 1.35 | .18 |

| Supplementary Table 2: Estimates of uncorrected GABA+ from Gannet 3.1 | | | | | | | | | | | | |
| --- | --- | --- | --- | --- | --- | --- | --- | --- | --- | --- | --- | --- |
|  | GABA+/ H_2_O | | | | GABA+/Cr | | | | GABA+/NAA | | | |
|  | Mean ± SD | | t | *p* | Mean ± SD | | t | *p* | Mean ± SD | | t | *p* |
|  | Young  adults | Older adults |  |  | Young  adults | Older  adults |  |  | Young adults | Older  adults |  |  |
| LPV | 1.85 ±0.16 | 1.75 ±0.23 | 2.74 | .007* | 0.14 ±0.01 | 0.13 ±0.02 | 4.46 | <.0001* | 0.08 ±0.01 | 0.08 ±0.01 | -0.98 | .33 |
| LVV | 1.60 ±0.20 | 1.38 ±0.24 | 5.68 | <.0001* | 0.13 ±0.02 | 0.11 ±0.01 | 5.39 | <.0001* | 0.07 ±0.01 | 0.07 ±0.01 | 1.18 | .24 |
| RVV | 1.74 ±0.22 | 1.51 ±0.23 | 5.97 | <.0001* | 0.13 ±0.01 | 0.12 ±0.01 | 6.99 | <.0001* | 0.08 ±0.01 | 0.07 ±0.01 | 2.53 | .01 |
| LAUD | 1.88 ±0.27 | 1.71 ±0.27 | 3.92 | .0001* | 0.12 ±0.02 | 0.11 ±0.02 | 3.46 | .0007* | 0.08 ±0.01 | 0.08 ±0.01 | 0.17 | .87 |
| RAUD | 1.89 ±0.19 | 1.70 ±0.25 | 4.69 | <.0001* | 0.13 ±0.01 | 0.12 ±0.01 | 5.88 | <.0001* | 0.09 ±0.01 | 0.08 ±0.01 | 1.53 | .13 |
| LSM | 2.10 ±0.32 | 1.91 ±0.27 | 3.79 | .0002* | 0.15 ±0.02 | 0.14 ±0.02 | 3.50 | .0006* | 0.08 ±0.01 | 0.08 ±0.01 | 1.11 | .27 |
| RSM | 2.17 ±0.21 | 1.98 ±0.22 | 5.10 | <.0001* | 0.16 ±0.02 | 0.15 ±0.02 | 3.52 | <.0006* | 0.08 ±0.01 | 0.08 ±0.01 | 0.63 | .53 |
| *survived correction for multiple comparison | | | | | | | | | | | | |

| Supplementary Table 3: Tissue corrected estimates of water referenced GABA+ from Gannet 3.1 | | | | | | | | |
| --- | --- | --- | --- | --- | --- | --- | --- | --- |
|  | GABA+/H_2_O CSF | | | | GABA+/ H_2_O TC | | | |
|  | Mean ± SD | | t | *p* | Mean ± SD | | t | *p* |
|  | Young adults | Older  adults |  |  | Young adults | Older  adults |  |  |
| LPV | 2.00 ±0.19 | 2.00 ±0.24 | -0.07 | .94 | 3.09 ±0.30 | 3.12 ±0.38 | -0.37 | .71 |
| LVV | 1.69 ±0.20 | 1.57 ±0.23 | 3.19 | .002* | 2.61 ±0.24 | 2.45 ±0.35 | 2.77 | .006* |
| RVV | 1.83 ±0.23 | 1.69 ±0.21 | 3.59 | .0004* | 2.81 ±0.30 | 2.63 ±0.32 | 3.21 | .001* |
| LAUD | 2.08 ±0.29 | 2.03 ±0.27 | 0.89 | .38 | 3.24 ±0.46 | 3.19 ±0.43 | 0.64 | .52 |
| RAUD | 2.14 ±0.19 | 2.08 ±0.25 | 1.49 | .14 | 3.35 ±0.30 | 3.28 ±0.39 | 1.21 | .23 |
| LSM | 2.30 ±0.34 | 2.27 ±0.25 | 0.69 | .49 | 3.57 ±0.53 | 3.55 ±0.39 | .25 | .80 |
| RSM | 2.43 ±0.23 | 2.41 ±0.27 | 0.82 | .42 | 3.80 ±0.36 | 3.78 ±0.45 | .28 | .78 |
|  | GABA+/ H_2_O ATC | | | | GABA+/ H_2_O GATC | | | |
|  | Mean ± SD | | t | *p* | Mean ± SD | | t | *p* |
|  | Young adults | Older  adults |  |  | Young adults | Older  adults |  |  |
| LPV | 2.92 ±0.27 | 3.00 ±0.36 | -1.38 | .17 | 2.22 ±0.21 | 2.28 ±0.27 | -1.38 | .17 |
| LVV | 2.54 ±0.30 | 2.41 ±0.34 | 2.42 | .02 | 1.85 ±0.22 | 1.75 ±0.24 | 2.42 | .02 |
| RVV | 2.73 ±0.34 | 2.58 ±0.33 | 2.62 | .01 | 2.00 ±0.25 | 1.89 ±0.24 | 2.63 | .01 |
| LAUD | 2.88 ±0.42 | 2.93 ±0.39 | -0.71 | .48 | 2.39 ±0.35 | 2.43 ±0.33 | -0.71 | .48 |
| RAUD | 2.95 ±0.28 | 2.97 ±0.35 | -0.45 | .65 | 2.47 ±0.23 | 2.49 ±0.29 | -0.45 | .65 |
| LSM | 3.51 ±0.51 | 3.63 ±0.40 | -1.36 | .17 | 2.52 ±0.36 | 2.59 ±0.28 | -1.36 | .17 |
| RSM | 3.70 ±0.35 | 3.81 ±0.45 | -1.54 | .13 | 2.70 ±0.25 | 2.74 ±0.32 | -1.54 | .13 |
| *survived correction for multiple comparisons | | | | | | | | |

| Supplementary Table 4: Estimates of GABA+/ H_2_O obtained using LCmodel | | | | |
| --- | --- | --- | --- | --- |
|  | Mean ± SD | | t | *p* |
|  | Young adults | Older adults |  |  |
| LPV | 2.76 ±0.18 | 2.67 ±0.31 | 1.90 | .06 |
| LVV | 2.41 ±0.28 | 2.11 ±0.35 | 5.43 | <.0001* |
| RVV | 2.61 ±0.28 | 2.26 ±0.34 | 6.65 | <.0001* |
| LAUD | 2.76 ±0.31 | 2.59 ±0.38 | 2.79 | .006* |
| RAUD | 2.81 ±0.24 | 2.60 ±0.33 | 4.17 | <.0001* |
| LSM | 2.50 ±0.26 | 2.31 ±0.30 | 3.88 | <.0001* |
| RSM | 2.61 ±0.29 | 2.45 ±0.38 | 2.73 | <.007 |
| *survived correction for multiple comparisons | | | | |
